# Single particle cryo-electron microscopy with an enhanced 200 kV cryo-TEM configuration achieves near-atomic resolution

**DOI:** 10.1101/2024.05.07.593029

**Authors:** Lijia Jia, Eliza A. Ruben, Humberto J. Suarez, Shaun K. Olsen, Elizabeth V. Wasmuth

## Abstract

Single particle cryogenic electron microscopy (cryo-EM) as a structural biology methodology has become increasingly attractive and accessible to investigators in both academia and industry as this ever-advancing technology enables successful structural determination of a wide range of protein and nucleic acid targets. Although data for many high resolution cryo-EM structures are still obtained using a 300 kV cryogenic transmission electron microscope (cryo-TEM), a modern 200 kV cryo-TEM equipped with an advanced direct electron detector and energy filter is a cost-effective choice for most single particle applications, routinely achieving sub 3 angstrom (Å) resolution. Here, we systematically evaluate performance of one such high-end configuration - a 200 kV Glacios microscope coupled with a Falcon 4 direct electron detector and Selectris energy filter (Glacios-F4-S). First, we evaluated data quality on the standard benchmarking sample, rabbit muscle aldolase, using three of the most frequently used cryo-EM data collection software: SerialEM, Leginon and EPU, and found that – despite sample heterogeneity – all final reconstructions yield same overall resolutions of 2.6 Å and map quality when using either of the three software. Furthermore, comparison between Glacios-F4-S and a 300 kV cryo-TEM (Titan Krios with Falcon 4) revealed nominal resolution differences in overall reconstructions of a reconstituted human nucleosome core particle, achieving 2.8 and 2.5 Å, respectively. Finally, we performed comparative data analysis on the human RAD51 paralog complex, BCDX2, a four-protein complex of approximately 150 kilodaltons, and found that a small dataset (≤1,000 micrographs) was sufficient to generate a 3.3 Å reconstruction, with sufficient detail to resolve co-bound ligands, AMP-PNP and Mg^+2^. In summary, this study provides evidence that the Glacios-F4-S operates equally well with all standard data collection software, and is sufficient to obtain high resolution structural information of novel macromolecular complexes, readily acquiring single particle data rivaling that of 300 kV cryo-TEMs.

## 1. Introduction

Cryo-electron microscopy (cryo-EM) has gained increased usage among a wide range of biologists investigating basic biological principles and molecular mechanisms to drug discovery due to its ability to resolve atomic details of vitrified particles in solution within hours. Resulting high resolution cryo-EM maps enable the building of atomic models for the purposes of discovery biology (for instance, architecture of novel complexes and active site chemistry), as well as structure-based drug design (Renaud *et al*., 2018; X. Zhang *et al*., 2021; Robertson, Meyerowitz and Skiniotis, 2022). In the past several years, recent advances in hardware and software have enabled structural determination of macromolecular machines at resolutions that exceed the theoretical Nyquist limit, with some systems reaching 1.2 Å resolution when imaged with a 300 kV cryo-TEM (Nakane *et al*., 2020; Yip *et al*., 2020; Zhang *et al*., 2020; Efremov and Stroobants, 2021; Feathers *et al*, 2021). With the integration of cryo-EM data collection and on-the-fly data processing software such as CryoSPARC Live (Punjani *et al*., 2017), obtaining a high resolution cryo-EM structure within one to two weeks is becoming more feasible (Baldwin *et al*., 2018; Efremov and Stroobants, 2021; Peck, Fay and Strauss, 2022). At present, there are two primary manufacturers of high end cryo-TEM instrumentation: ThermoFisher Scientific and Jeol, both of whom offer high acceleration voltage high-throughput 300 kV cryo-TEMs for single particle and cryo-electron tomography applications, including the Titan Krios and CRYO ARM 300, respectively. These manufacturers also offer lower acceleration voltage 200 kV cryo-TEMs that historically have been considered screening microscopes for single particle cryo-EM, including the Jeol CRYO ARM 200 (Merk *et al*., 2020), ThermoFisher’s Talos Arctica (Herzik, Wu and Lander, 2017; Cash *et al*., 2020; Peck, Fay and Strauss, 2022) and ThermoFisher’s Glacios (Hamdi *et al*., 2020).

However, recent technological advances demonstrate that when equipped with state-of-the-art hardware, including a direct electron detector (such as the Falcon 4/4i) and a high quality post-column energy filter to efficiently remove background from inelastically scattered electrons (such as the Selectris and Selectris X), and when housed in an ideal environment (optimal temperature, humidity, and minimal vibration and electromagnetic interference), high-resolution and high-contrast imaging is enabled on 200 kV cryo-TEMs, permitting acquisition of near atomic resolution structural models by single particle cryo-EM (Nakane *et al*., 2020; Keizer *et al*., 2021; Kern *et al*., 2021; Fréchin *et al*., 2022; Adrian *et al*., 2022). As a result, ThermoFisher Scientific’s Glacios 200 kV cryo-TEM has emerged as an attractive option for researchers interested in obtaining high-resolution structures of biological macromolecules but also seeking lower-cost and smaller footprint solutions compared to ThermoFisher’s other 200 kV and 300 kV cryo-TEMs.

The Glacios 200 kV microscope is often integrated with ThermoFisher Scientific’s EPU data collection software, a license-based software that boasts a simple, user-friendly graphical user interface, including capabilities for autoalignments, automated data acquisition, ice quality filters, multi-grid data collection, and on-the-fly image analysis through Smart EPU (Sader *et al*., 2020; Kimanius *et al*., 2021; Ye, Liu and Li, 2022).

In addition to EPU, Leginon and SerialEM are two widely used open-source data collection software. Leginon was developed by the National Resource for Automated Molecular Microscopy (NRAMM) at Scripps Research Institute (Potter *et al*., 1999; Carragher *et al*., 2000; Cheng *et al*., 2021), and has a strong user base with ongoing development. It features a modular design that allows for easy customization and integration with other software tools, modalities to measure ice thickness and other grid quality metrics, has its own web server and database providing remote collaborations and live analysis and feedback, and offers a wide range of data collection protocols, including single-particle analysis, cryo-electron tomography, and cryo-electron diffraction (Cheng *et al*., 2023). SerialEM is a script-based software developed by the Boulder Laboratory for 3D Electron Microscopy of Cells at the University of Colorado (Mastronarde, 2003). This software also features ongoing development and support. It is designed for high-throughput data collection, and is ideally suited for automated acquisition of tilt series for cryo-electron tomography (Mastronarde, 2005; Burt *et al*., 2021). Its strengths include versatility, compatibility with multiple hardware platforms, powerful features for tomography, and advanced montaging and autofocus capabilities. While both Leginon and SerialEM are freely available, they are less intuitive to use as compared to EPU, and thus require more specialized training to operate.

Here, we aim to systematically assess the performance of a high end 200 kV microscope, a Glacios configured with a Falcon 4 detector and Selectris energy filter (henceforth referred to as Glacios-F4-S). We begin by benchmarking performance, data quality, and resolution of resulting 3D reconstructions when operating the Glacios with the three most popular data acquisition software packages, EPU, Leginon, and SerialEM, on a standard benchmarking sample, rabbit muscle aldolase. We next compared performance of the Glacios-F4-S to a 300 kV Titan Krios equipped with a Falcon 4 in resolving the structure of a human reconstituted nucleosome core particle. Finally, we demonstrate that as few as 1,000 micrographs (5 hours of data collection) are sufficient for *de novo* structural resolution of a small (<150 kDa) human DNA repair complex, BCDX2, and co-bound AMP-PNP and magnesium ligands.

## 2. Methods

### 2.1. Sample preparation

Aldolase purified from rabbit muscle was purchased from Sigma-Aldrich (catalogue number A2714, ≥80% biuret) in the form of lyophilized powder and dissolved to a concentration of 1 mg/mL in 20 mM HEPES PH 7.5 and 50 mM NaCl. To purposefully maintain sample heterogeneity, no further purification by size exclusion chromatography was performed, bypassing the standard protocol for structural determination of this cryo-EM benchmark sample (Herzik *et al*., 2017). To assess the oligomeric composition of the sample, mass photometry analysis of resuspended aldolase was carried out using a Refeyn TwoMP instrument. Contrast-to-mass calibration was achieved by a standard curve covering a mass range from 66 up to 480 kDa. The experiments were carried out on sample carrier slides partitioned by six-well silicone cassettes (Refeyn). Using AcquireMP software, a blank acquisition field was first adjusted in native settings using sterile-filtered PBS buffer, and then the 1 mg/mL resuspended aldolase stock was flash diluted into PBS buffer to a final concentration of 12 nM. Video recordings were immediately performed to detect single particles over the course of 1 min at a rate of 100 frames per second in ratiometric acquisition settings. All mass photometry data were analyzed with DiscoverMP software to produce mass values for each detected particle and plotted as normalized counts. Aldolase sample purity was further assessed by SDS-PAGE and stained with Coomassie.

For human nucleosome assembly, standard protocols were used as described by Dyer *et al*., 2003, with some modifications. Briefly, bacterial pRSF-Duet1 plasmids harboring H2A-H2B or H3-H4 of the human nucleosome were expressed in *E. coli* expression strain BL21 DE3 codon plus (RILP). Cell pellets were resuspended in lysis buffer (20 mM Tris-HCl pH7.8, 300 mM NaCl, 1 mM DTT and 1 mM EDTA) and lysed by sonication. Lysates were clarified by high speed centrifugation, and the H2A-H2B dimer and H3-H4 tetramer were subsequently purified by heparin affinity chromatography followed by size exclusion chromatography (HiLoad 26/600 Superdex 200 column, Cytiva Life Sciences). To prepare the histone octamer, H2A-H2B and H3-H4 at a 1:2 molecular ratio were mixed together at 20 micromolar concentration in 20 mM Tris-HCl pH 7.8, 500 mM NaCl, 1 mM DTT, 1 mM EDTA and 7 M guanidine hydrochloride. After a 2 hour incubation, the mixture was dialyzed into gel filtration buffer (20 mM Tris-HCl pH7.8, 2 M NaCl, 1 mM DTT and 1 mM EDTA) and purified by size exclusion chromatography on a HiLoad 26/600 Superdex 200 column (Cytiva Life Sciences). A relative histone octamer peak was apparent, and fractions collected. 187 base pairs of nucleosome core particle (NCP) DNA was amplified by PCR from the plasmid pBluescript II SK (+) encoding NCP DNA and further purified by ion exchange chromatography on a MonoQ 10/100 column (Cytiva Life Sciences).

Nucleosomes were reconstituted by sequential dialysis where histone octamers were mixed with nucleosome core particle DNA at a molar ratio of 1:1.2 beginning at 20 mM Tris-HCl pH 7.8, 2 M KCl, 1 mM DTT and 1 mM EDTA dialysis and ending at 250 mM KCl over three rounds of overnight dialysis. Nucleosomes were further purified by running the reconstituted complex through a DEAE anion exchange column (TSKgel DEAE-5PW, Tosoh Biosciences). Peaks corresponding to intact nucleosomes were collected, concentrated to 0.3 mg/mL, and used for grid preparation.

Recombinant human BCDX2 was purified from insect cells and reconstituted as described previously (Rawal *et al*., 2023).

### 2.2 Grid vitrification

For cryo-EM grid preparation, all grids were glow-discharged in air at 20 mA for 30 seconds before applying sample. 3 ul mouse apoferritin sample was applied onto a Quantifoil Cu holey carbon grid (R2/2, 200 mesh); 3 ul rabbit aldolase, human nucleosome, or human BCDX2 was applied onto UltrAuFoil grid (R1.2/1.3, 300 mesh). The samples were vitrified at 8 °C and 100% humidity using a ThermoFisher Scientific Vitrobot Mark IV by blotting for 3 s, with a blot force of 0. Grids were plunge frozen in liquid ethane and transferred into liquid nitrogen for storage.

### 2.3. Cryo-EM data acquisition

Unless otherwise noted, all data were collected on the Glacios-F4-S at the University of Texas Health Sciences Center at San Antonio (UTHSCSA). For data collection on the Glacios-F4-S, a cross-grating grid was loaded into the microscope and proper eucentric height determined using a spot size of 4 and a 50 uM C2 aperture to perform initial beam alignment. For data acquisition, the eucentric focus was manually set and diffraction lens turned on to diffraction mode, followed by inserting the 100 uM objective aperture and adjusting the beam intensity to ensure a sharp spread of diffraction bands. A 130K magnification with a pixel size 0.87 Å was used for all subsequent data collection (aldolase, nucleosome, and BCDX2). With EPU, the embedded AFIS model was used for data collection. In order to use the aberration correction model during data collection in Leginon and SerialEM for aldolase, image-shift, astigmatism and coma-free calibration were performed at a magnification of 130K before data acquisition. A 3X3 hole image shift model was used for the data collection. The defocus range used was from −0.8 to −2.0. A total of 1,1741 image frames with EER file format were recorded with the Falcon 4 camera, with an exposure time of 7 s, with a 5.65 ē/pix/s dose rate, for a total dose of 54 ē/Å^2^.

Nucleosome data were collected on a 300 kV ThermoFisher Scientific Titan Krios G4 microscope with a Falcon 4 detector at the National Center for Cryo-EM Access and Training (NCCAT) at a pixel size of 0.833 Å and a total dose of 56 ē/Å^2^, in addition to the dataset acquired at 200 kV on the Glacios-F4-S at UTHSCSA at a pixel size of 0.87 Å and at a total dose of 54 ē/Å^2^.

### 2.3 Cryo-EM data processing

Image frames were gain corrected, dose weighted, patch motion corrected using CryoSPARC Live v3.3.1 during data collection (Punjani *et al*., 2017). The exact defocus value and CTF (Contrast Transfer Function) of each micrograph was estimated using CryoSPARC’s patch CTF estimation. Micrograph curation with a CTF resolution cutoff value of 5 Å resolution was applied to all data. Automated blob picking was used exclusively for aldolase and BCDX2 datasets. For the nucleosome, the blob picker function in CryoSPARC was first used for initial automated picking, yielding featured 2D classes that were consequently used for template picking. Particle stacks for all three complexes were subsequently extracted at a 256 pixel box size without downsampling for aldolase, nucleosome, and BCDX2. After three rounds of light 2D classification to filter noise and dissociated particles, the resulting particle stacks were input for the next stages of reference-free 3D ab initio reconstruction, followed a round of 3D heterogeneous refinement, the exception being BCDX2 which underwent several rounds of heterogenous refinement as described previously (Rawal *et al*., 2023). The best classes were used for homogeneous refinement, applying D2 symmetry for aldolase, and C1 for the human nucleosome and BCDX2.

### 2.4 Cryo-EM map model building and refinement

Models of aldolase and the nucleosome were obtained from the protein data bank (RCSB PBD), and coordinates of BCDX2 were obtained from our previous work (Rawal *et al*., 2023). Rigid-body docking into the cryo-EM map was done in UCSF ChimeraX (Pettersen *et al*., 2021). Further building and structural refinements were performed using COOT (Emsley *et.al.*, 2004) and further refinements of stereochemistry and secondary structure were performed in PHENIX using phenix.real_space_refine (Liebschner *et.al.*, 2019). All coordinates were validated using wwPDB (https://validate-rcsb-2.wwpdb.org/). Figures for the cryo-EM map and models were prepared with ChimeraX and PyMOL.

## 3. Results

### 3.1. Aldolase data collection on the Glacios-F4-S: a comparison of data acquisition software on a heterogenous sample

Aldolase is a metabolic enzyme that forms a 170 kDa tetramer compromised of four 42.5 kDa monomers. The D2 symmetry of the tetramer form makes it amenable for benchmarking performance of a wide range of cryo-TEMs (Kim *et al*., 2018; Li *et al*., 2020; Wu, Lander and Herzik, 2020). To assess the performance of the Glacios-F4-S on heterogenous data, we vitrified aldolase isolated from rabbit muscle that did not undergo additional purification by gel filtration to purposely maintain a mixture of monomer, dimer, and higher oligomer populations as well as higher molecular weight contaminants, as evidenced by mass photometry and SDS-PAGE (**Fig. S1a,b**), as an increasing number of single particle cryo-EM projects involve a significant degree of intrinsic compositional and conformational heterogeneity.

We collected three datasets using EPU, SerialEM, and Leginon from a single grid. Notably, micrographs were acquired at a rate of 180 micrographs per hour for EPU, and 140 micrographs per hour for SerialEM and Leginon (see more in Discussion). Datasets were comprised of 3,117 micrographs from EPU, 3,600 micrographs from SerialEM, and 3,228 micrographs from Leginon and were subject to further data processing and comparison. After a series of processing steps in CryoSPARC, including automated particle picking, 2D classification, reference-free ab-initio reconstruction, and heterogenous refinement, finalized clean particle stacks of 111,600 (EPU), 111,059 (SerialEM), and 111,500 (Leginon) particles underwent homogenous refinement with C1 and D2 symmetry applied (**Table 1**). Despite collecting from a preparation of aldolase that is 33% tetramer and is comprised primarily of a mixture monomers, dimers, and higher order oligomers (**Fig. S1a,b**), the final overall resolution and quality of the resulting tetramer reconstructions were high and comparable, with EPU yielding a 2.6 Å model with the lowest B factor of −87.8 Å^2^, SerialEM yielding a 2.7 Å with a B factor of −93.5 Å^2^, and Leginon yielding a 2.6 Å model with a B factor of −94.2 Å^2^ (**Fig. 1a-c, Fig. S1c-e**). From the standpoint of local resolution, all three reconstructions are similar, with the highest resolution in the central core (2.5 Å) and lower resolution in the peripheral surface (3.0 Å). Further atomic model fitting and model building also demonstrated no obvious differences in map quality in data collected with EPU, SerialEM or Leginon, as amino acid side chain density is prominent and highly featured in all three datasets (**Fig. 1d,e**).

**Figure 1.**
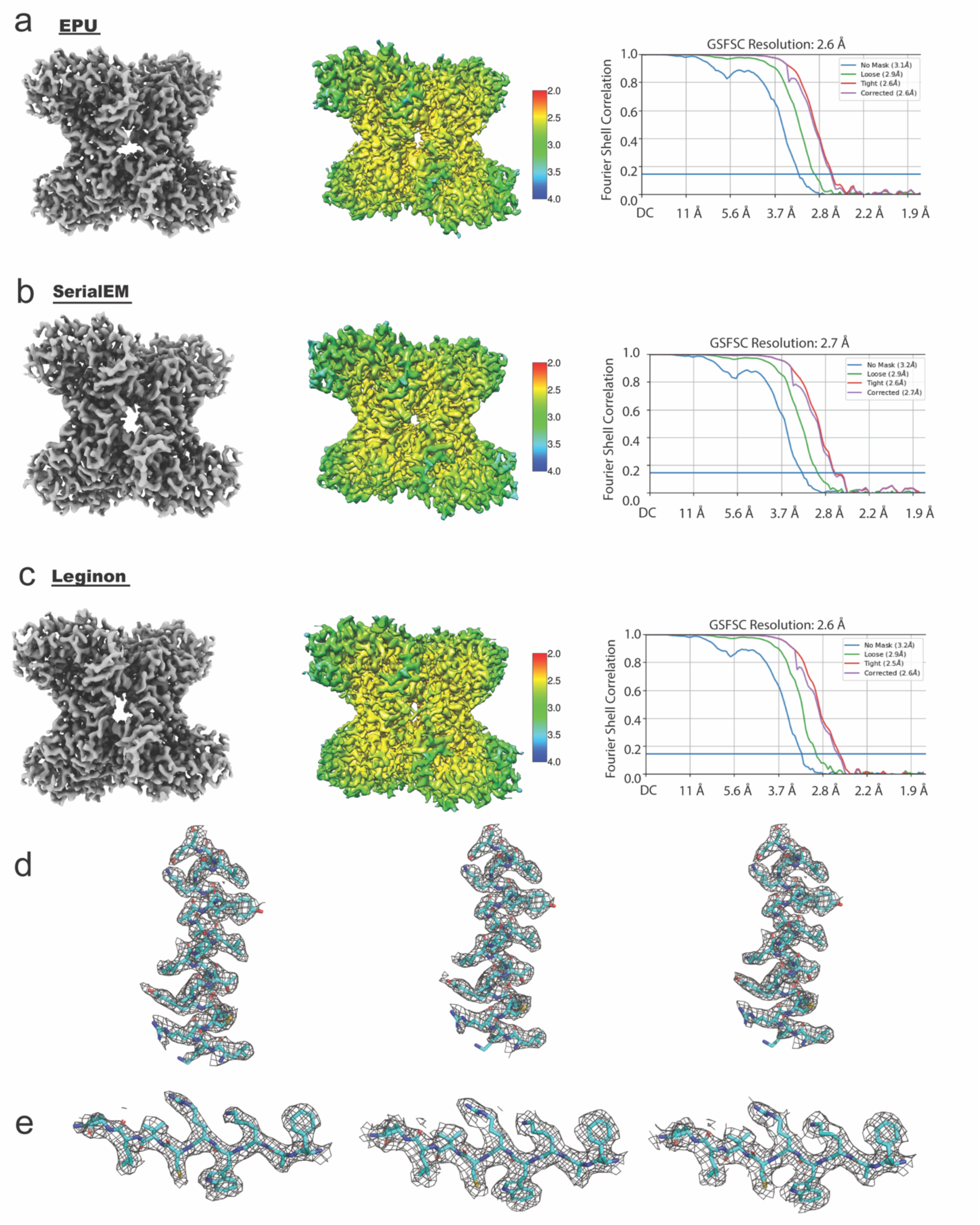
Rabbit muscle aldolase cryo-EM reconstructions obtained at 200 kV with three data collection software. (**a**) Final reconstructed map from data collected using EPU software (left). Local resolution map (middle) color-coded from lower (blue) to higher resolution (red) indicates a resolution range between 2.5-3.0 Å. The Fourier Shell Correlation (FSC) plot from two half maps reveals an overall resolution of 2.6 Å as determined at 0.143 criterion (right). (**b**) Final reconstructed map from data collected using SerialEM software (left). Local resolution map (middle) indicates a resolution range between 2.5-3.0 Å. The FSC plot from two half maps reveals an overall resolution of 2.7 Å as determined at 0.143 criterion (right). (**c**) Final reconstructed map from the data collected using Leginon software (left). Local resolution map (middle) indicates a resolution range between 2.5-3.0 Å. The FSC plot from two half maps reveals an overall resolution of 2.6 Å as determined at 0.143 criterion (right). (**d,e**) Local electron density maps of side chains 198-218 (**d**) and 144-152 (**e**) from EPU (left), SerialEM (middle), and Leginon (right) respectively.

**Table 1:**
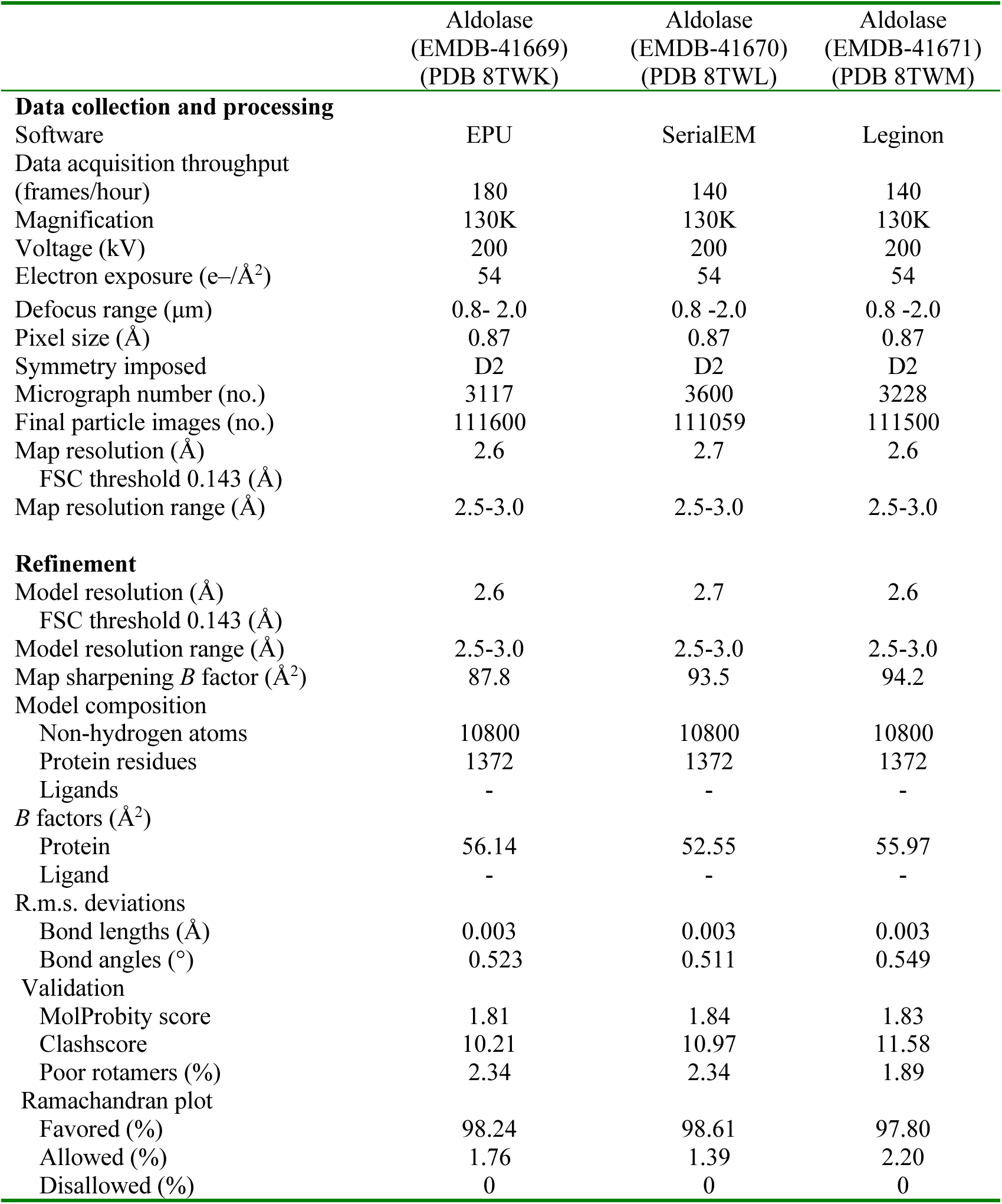
Cryo-EM data collection, refinement and validation statistics for aldolase.

Of note, the structure of tetrameric aldolase has previously been resolved to 2.1 Å at 200 kV (Wu *et al*., 2020). This dataset was collected at higher magnification (0.56 Å/pixel versus 0.87 Å/pixel) and was a larger dataset (3,534 micrographs, 1.8 million particles) comprised of highly pure aldolase tetramer that had been enriched before imaging by size exclusion chromatography. While our reconstruction is lower resolution (2.6 Å), the dataset was smaller, with 1.1 million particles, of which approximately a third or less were the intact tetramer.

### 3.2. Nucleosome core particle data collection and analysis at 200 and 300 kV

The human nucleosome core particle is a non-symmetrical nucleoprotein complex comprised of high affinity double-stranded DNA interactions wrapped around a histone octamer that form the basic unit of chromatin. High resolution (<4 Å) structures of reconstituted mononucleosomes have historically been collected at 300 kV. To evaluate whether the Glacios-F4-S can acquire high resolution 3D reconstructions of a human nucleosome comparable to that obtained on a Titan Krios G4 with a Falcon 4, we collected two small datasets on two replicate grids, with 1,500 micrographs collected on the Titan Krios (300 kV) at a total dose of 56 e-/Å^2^, and 1,300 micrographs collected on the Glacios-F4-S (200 kV) at a total dose of 54 e-/Å^2^. A general data processing workflow in CryoSPARC was subsequently implemented, including creation of 2D templates for template-based particle picking, 2D classification of the entire dataset, followed by reference-free ab-initio reconstruction and heterogenous refinement. A final particle stack of 174,000 particles (Titan Krios) and 174,571 particles (Glacios-F4-S) were used for the final steps of homogenous refinement and non-uniform refinement (**Table 2**). Interestingly, differences in overall resolution were minimal, with the final resolution of 3.0 Å with a relative B factor of −115 Å^2^ for the Krios dataset, and 3.3 Å with a relative B factor of −97.3 Å^2^ for the Glacios dataset (**Fig. 2a,b**). To our knowledge is the highest resolution structure of a human nucleosome obtained at 200 kV.

**Figure 2.**
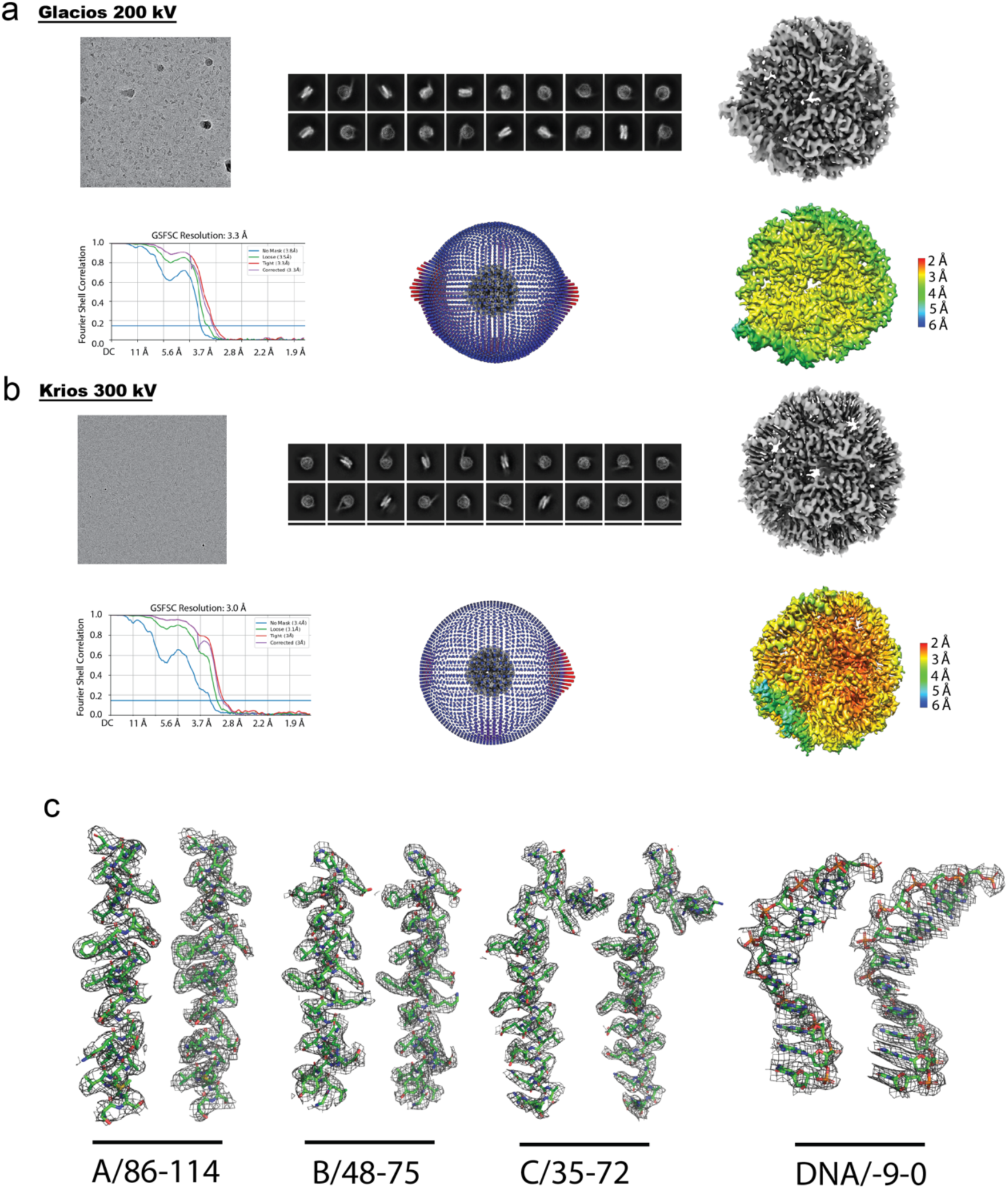
Comparison of human nucleosome data quality and reconstructions collected at 200 kV (Glacios) and 300 kV (Krios) cryo-TEMs equipped with Falcon 4 detectors. (**a**) Representative micrograph imaged at 200 kV with a Glacios cryo-TEM equipped with Selectris energy filter (top left), 2D classification (top middle), and density map after data processing (top right), Fourier Shell Correlation (FSC) plot from two half maps, with an overall resolution of 3.3 Å as evaluated at 0.143 criterion (bottom left), Euler angle distribution of particles used for 3D homogenous refinement (bottom middle), and local resolution map color-coded from lower (red) to higher resolution (blue) (bottom right). (**b**) As in (a) except data were collected at 300 kV on a Titan Krios cryo-TEM. (**c**) Representative side chains and nucleotides built into local density maps, with data derived from Glacios on the left and Krios on the right. Histone H3.2 (Chain A) residues shown from 86 to 114. Histone H4 (Chain B) residues shown from 48 to 75. Histone H2A (Chain C) residues shown from 35 to 72 and DNA nucleotides −9 to 0 shown.

**Table 2:** Cryo-EM data collection, refinement and validation statistics for nucleosome.

|  | Nucleosome<br>(EMDB-44470)<br>(PDB 9BE5) | Nucleosome<br>(EMDB-44471)<br>(PDB 9BE6) |
| --- | --- | --- |
| **Data collection and processing** |  |  |
| Microscope | Glacios | Krios |
| Data collection software | EPU | Leginon |
| Data acquisition throughput<br>(frames/hour) | 180 | 145 |
| Magnification | 130K | 96K |
| Voltage (kV) | 200 | 300 |
| Electron exposure (e-/Å<sup>2</sup>) | 54 | 56 |
| Defocus range (µm) | 0.8 - 2.0 | 0.8 - 2.0 |
| Pixel size (Å) | 0.87 | 0.833 |
| Symmetry imposed | C1 | C1 |
| Micrograph number (no.) | 1300 | 1500 |
| Final particle images (no.) | 174571 | 174000 |
| Map resolution (Å) | 3.3 | 3.0 |
| FSC threshold 0.143 (Å) |  |  |
| Map resolution range (Å) | 2.8-3.6 | 2.5-3.8 |
| **Refinement** |  |  |
| Model resolution (Å) | 3.3 | 3.0 |
| FSC threshold 0.143 (Å) |  |  |
| Model resolution range (Å) | 2.8-3.6 | 2.5-3.8 |
| Map sharpening *B* factor (Å<sup>2</sup>) | 104.4 | 129.3 |
| Model composition |  |  |
| Non-hydrogen atoms | 11898 | 11163 |
| Protein residues | 751 | 751 |
| Nucleotide | 290 | 254 |
| *B* factors (Å<sup>2</sup>) |  |  |
| Protein | 47.38 | 116.31 |
| Nucleotide | 104.37 | 195.13 |
| R.m.s. deviations |  |  |
| Bond lengths (Å) | 0.004 | 0.005 |
| Bond angles (°) | 0.581 | 0.575 |
| Validation |  |  |
| MolProbity score | 1.32 | 1.01 |
| Clashscore | 5.72 | 2.29 |
| Poor rotamers (%) | 0 | 0 |
| Ramachandran plot |  |  |
| Favored (%) | 97.96 | 98.23 |
| Allowed (%) | 2.04 | 1.77 |
| Disallowed (%) | 0 | 0 |

Although differences in overall resolution were minimal, more notable differences were observed at the level of local resolution (**Fig. 2a,b**), with maps obtained on the Titan Krios showing a resolution range from 2.2 to 3.2 Å, and Glacios-F4-S maps covering a range from 3.0 to 4.0 Å. While clear amino acid side chain density within each of the subunits of the histone octamer was evident and comparable in both maps, electron density for the co-bound double-stranded DNA was noticeably weaker in data obtained on the Glacios-F4-S compared to the Titan Krios, with a resolution difference of approximately 1 Å (**Fig. 2c**). Potential rationale for the disparity in nucleic acid density between the two datasets may be due to decreased signal to noise ratio of solvent-exposed and more conformationally dynamic entities on the periphery of the complex at 200 kV compared to 300 kV, and/or due to the increased probability of ionization events at lower acceleration voltages (Grubb and Groves, 1971). Future experiments investigating how dataset size and total dose address the disparity in nucleic acid electron density between nucleosome datasets obtained on the Glacios-F4-S and Titan Krios (**Fig. 2c**) will be of interest to investigators in the cryo-EM community studying structure of nucleic acids and nucleoprotein complexes.

### 3.3. The Glacios-F4-S is sufficient to acquire high resolution data of the human RAD51 paralogue complex, BCDX2

BCDX2 is a tetrameric complex comprised of the RAD51 paralogues RAD51B, RAD51C, RAD51D, and XRCC2. Despite its essential role in homologous recombination and genome stability and its frequent mutation in cancer, fundamental questions pertaining to architecture, ligand binding, and DNA substrate recognition were unknown due to its recalcitrance to crystallization, relatively small size (150 kDa) and lack of symmetry, making it a historically challenging sample for cryo-EM and X-ray crystallography. We recently described some of the first high resolution cryo-EM reconstructions of BCDX2 in the absence and presence of single-stranded DNA, resolved to 2.3 and 3.1 Å, respectively (Rawal *et al*., 2023), The structures revealed how the complex is assembled and provide insight to nucleotide binding within each of the subunit’s conserved ATPase domains, as well as the molecular determinants of DNA substrate specificity. Notably, these data were exclusively acquired on the Glacios-F4-S, a remarkable technological achievement given the small size of the complex (with 97 kDa of 150 kDa of the complex ordered in our structural models), and the relatively modest size of the datasets. Approximately 5,600 micrographs for the apo dataset were sufficient to obtain a 2.3 Å reconstruction consisting of 489,073 particles, and approximately 4,000 micrographs for the single stranded DNA-bound dataset resulted in a 3.1 Å consisting of 165,968 particles with unambiguous density for DNA (Rawal *et al*., 2023).

To determine the minimal amount of data required for *de novo* structure determination of the complex, as well as visualization of nucleotide binding, we iteratively processed 1,000, 1,500, and 5,000 micrographs of BCDX2 co-bound to the non-hydrolyzable ATP analogue, AMP-PNP, and its cofactor, Mg^+2^ (**Fig. 3, Fig. S3**). At a collection rate of 180 micrographs per hour, this equates to 5.5, 8.3, and 28 hours of data collection. Within all three sets of micrographs, clear secondary structure features are apparent in calculated 2D classes (**Fig. 3a**). Similarly, all three sets of micrographs yielded high resolution reconstructions, with even the small dataset (1,000 micrographs, 71,500 particles) reaching an overall resolution of 3.1 Å, and showing contiguous density for co-bound AMP-PNP and Mg^+2^ (**Fig. 3a**, **Table 3**). Map quality and overall resolution continued to improve non-linearly with additional data, with the 1,500 micrograph dataset reaching an overall resolution of 2.9 Å and consisting of 113,054 particles (**Fig. 3b**), and the 5,000 micrograph dataset reaching an overall resolution of 2.3 Å and consisting of 375,146 particles (**Fig. 3c**).

**Figure 3.**
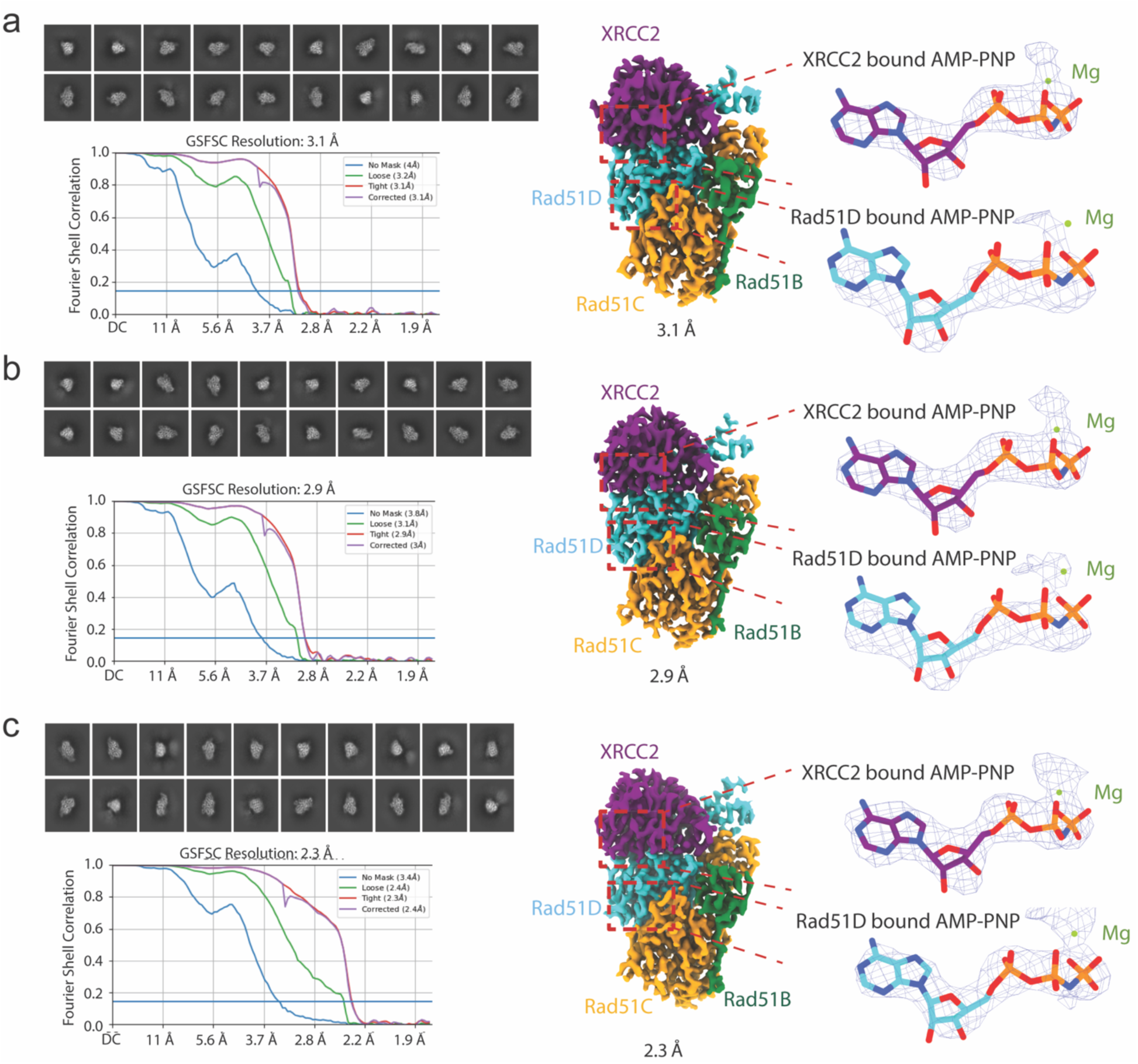
Comparison of human BCDX2 overall reconstructions and electron density collected at 200 kV (Glacios-F4-S). (**a-c**) Representative 2D classification (upper left), Fourier Shell Correlation (FSC) plots (lower left), and electron density maps (right) from 1,000 (**a**), 1,498 (**b**), and 5,000 (**c**) micrographs of human reconstituted BCDX2 images on the Glacios-F4-S. Zoom insets in electron density maps highlight two AMP-PNP molecules and co-bound magnesium ions.

**Table 3:** Cryo-EM data collection, refinement and validation statistics for BCDX2.

|  | BCDX2 | BCDX2 | BCDX2 |
| --- | --- | --- | --- |
| **Data collection and processing** |  |  |  |
| Microscope | Glacios | Glacios | Glacios |
| Data collection software | EPU | EPU | EPU |
| Data acquisition throughput (frames/hour) | 180 | 180 | 180 |
| Magnification | 130K | 130K | 130K |
| Voltage (kV) | 200 | 200 | 200 |
| Electron exposure (e-/Å<sup>2</sup>) | 51 | 51 | 51 |
| Defocus range (µm) | 0.8 -2.0 | 0.8 -2.0 | 0.8 -2.0 |
| Pixel size (Å) | 0.87 | 0.87 | 0.87 |
| Symmetry imposed | C1 | C1 | C1 |
| Micrograph number (no.) | 1000 | 1498 | 5000 |
| Final particle images (no.) | 71500 | 113054 | 375146 |
| Map resolution (Å) | 3.12 | 2.95 | 2.35 |
| FSC threshold 0.143 (Å) |  |  |  |
| Map resolution range (Å) | 2.8-5.0 | 2.5-5.0 | 2.2-4.5 |
| **Refinement** |  |  |  |
| Model resolution (Å) | 3.12 | 2.95 | 2.35 |
| FSC threshold 0.143 (Å) |  |  |  |
| Model resolution range (Å) | 2.9-3.9 | 2.6-3.6 | 2.6-3.4 |
| Map sharpening *B* factor (Å<sup>2</sup>) | 63.8 | 68.5 | 54.7 |
| Model composition |  |  |  |
| Non-hydrogen atoms |  |  |  |
| Protein residues | 7280 | 7280 | 7280 |
| Ligands | 91 | 91 | 91 |
| *B* factors (Å<sup>2</sup>) |  |  |  |
| Protein | 67.8 | 67.8 | 67.8 |
| Ligands | 61.8 | 61.8 | 61.8 |
| R.m.s. deviations |  |  |  |
| Bond lengths (Å) | 0.002 | 0.002 | 0.002 |
| Bond angles (°) | 0.444 | 0.444 | 0.444 |
| Validation |  |  |  |
| MolProbity score | 1.6 | 1.6 | 1.6 |
| Clashscore | 10.4 | 10.4 | 10.4 |
| Poor rotamers (%) | 1.98 | 1.98 | 1.98 |
| Ramachandran plot |  |  |  |
| Favored (%) | 97.6 | 97.6 | 97.6 |
| Allowed (%) | 2.4 | 2.4 | 2.4 |
| Disallowed (%) | 0 | 0 | 0 |

Concurrent with our initial report of the apo and ssDNA-bound BCDX2 structures, Greenhough and colleagues reported similar structures of BCDX2 obtained from a larger, 35,000 micrograph dataset on a Titan Krios equipped with a K3 detector (Greenhough *et. al*, 2023). While we cannot exclude contributions of sample and grid preparation, as well as differences in reconstruction workflows in accounting for these differences, it is important to note that final resolutions and atomic details were comparable in both studies.

Overall, these data suggest that the Glacios-F4-S is sufficient for *de novo*, high resolution structure determination of a small, structurally elusive and important DNA repair complex, generating data sufficient to reveal previously unknown architecture as well as co-bound AMP-PNP/Mg^+2^ in 5 hours, mirroring results obtained at 300 kV.

## 4. Discussion

### 4.1 Summary of findings

The cryo-EM resolution revolution and continuing technological advances in the field have made the methodology a first-line choice for an increasing number of structural biology applications, from discovery of overall architecture to resolution of small molecule binding pockets. With rapid and continued development of hardware, data collection software, and post-processing algorithms, infrastructure-associated hurdles serve as the more frequently encountered bottlenecks for successful structure determination. These include regular access to high-end and high-throughput instrumentation, as well as the high costs associated with cryo-TEM purchase, maintenance, and user fees. Although cryo-EM access is available free of cost at several national centers, users typically encounter several month wait times before a scheduled session and may need to submit peer-reviewed proposals with preliminary data demonstrating sample readiness. Academic facilities may also accessible, but may charge higher rates to external users, who may also experience an extended wait time depending on the needs of the facility’s internal user base.

The study presented here showcases the versatility of the Glacios-F4-S in resolving high resolution structures of a range of molecular targets, including a heterogenous protein preparation of aldolase, and two non-symmetric human DNA binding complexes – the nucleosome and BCDX2. We demonstrate that this configuration can resolve ligands, including nucleic acid, nucleotides, and even metal cofactors, with several of the maps rivaling that obtained at 300 kV with state-of-the-art direct electron detectors (Falcon 4 and K3). Given that the cost of a Glacios-F4-S is significantly less than a Titan Krios in terms of housing requirements, upfront instrument costs, and annual service contract, our conclusion is that this 200 kV setup may very likely meet the needs of most single particle cryo-EM investigators, particularly in situations where a 300 kV cryo-TEM is cost-prohibitive. Steps should be taken to ensure that the specimens imaged are comprised of vitreous ice thin enough to yield high particle contrast while minimizing background noise at 200 kV, yet thick enough to yield sufficient particle density, maximize particle views, and minimize sample dissociation to yield a high resolution 3D reconstruction.

Furthermore, our study demonstrates that data generated from the Glacios-F4-S is comparable among all three of the commonly used data collection software: EPU, SerialEM, and Leginon. However, while our study required that we compare an equal number of particles for analysis, there was an appreciable decrease in throughput with the two open-source software. As SerialEM and Leginon are installed on separate computers, delays in data collection are introduced as commands are transferred from the computer in which SerialEM or Leginon are installed to the computer controlling the microscope. In contrast, EPU is installed on the computer controlling the microscope. Furthermore, EPU applies a larger image-shift range of 5-9 μm, while SerialEM and Leginon used a 3.5 μm image-shift, resulting in increased stage movement deployed by the latter two software. Taken together, these differences account for nearly a 1,000 micrograph difference in a 24 hour period between EPU and SerialEM/Leginon. In the example with aldolase, EPU collected at a rate of 180 micrographs per hour (or 4,320 micrographs in a 24 hour period) while Leginon and SerialEM collected at a rate of 140 micrographs per hour (or 3,360 micrographs in a 24 hour period).

### 4.2 Future perspectives

Lastly, as hardware and software upgrades designed to increase the rate of data collection from individual foil holes continue to be developed, relative differences in performance between the 200 kV configuration described and that of high-end 300 kV cryo–TEMs are minimized for single particle specimens with high quality vitreous ice. At the time of writing of this manuscript, over 75% of all EMDB entries were generated at 300 kV and approximately 12% of entries were generated at 200 kV. After completion of this study, recent Glacios-specific developments released from ThermoFisher include implementation of a cold-FEG, upgrades for a faster camera (Falcon 4i), as well as upgrades for aberration-free image shift (AFIS), and fringe free imaging (FFI) eucentric, which enable the user to acquire multiple shots per foil hole. When combined, the Falcon 4i and FFI upgrades can substantially increase throughput, from 5,000 micrographs per 24 hour period to up to nearly 20,000 micrographs per 24 hour period, addressing a major bottleneck in data acquisition at 200 kV compared to 300 kV. Future studies will reveal how this enhanced throughput increases the likelihood of solving the most challenging single particle cryo-EM projects at 200 kV.

## Acknowledgements

The 200 kV cryo-EM data used for all projects were collected at the UT Health San Antonio Cryo-EM Facility on a Glacios transmission electron microscope equipped with a Falcon 4 camera and a Selectris energy filter purchased with the support of University of Texas STARs award 402-1317 (S.K.O.). Some of this work was performed at the National Center for Cryo-EM Access and Training (NCCAT) and the Simons Electron Microscopy Center located at the New York Structural Biology Center, supported by the NIH Common Fund Transformative High Resolution Cryo-Electron Microscopy program (U24 GM129539) and by grants from the Simons Foundation (SF349247) and NY State Assembly Majority. We thank Edward Eng for his insights.

Research reported in this publication was supported by NIH grants R01 GM115568 and R01 GM128731 (S.K.O.) and R00 GM140264 (E.V.W.). S.K.O. is the recipient of a Cancer Prevention and Research Institute of Texas (CPRIT) Rising Star Award (RR200030), and E.V.W. is the recipient of a Cancer Prevention and Research Institute of Texas (CPRIT) Recruitment of First Time Tenure Track Faculty Award (RR220068). This research was also supported in part by the Prostate Cancer Foundation Young Investigator Award (E.V.W.) and the Voelcker Foundation Young Investigator Award (E.V.W.).

## Author Contributions

E.V.W. conceived of the study. L.J. and E.V.W. designed experiments. L.J. and H.J.S. performed experiments. L.J., E.A.R., S.K.O. and E.V.W. analyzed data and wrote the manuscript. All authors edited the manuscript.

## References

Baldwin, P.R. et al. (2018) ‘Big data in cryoEM: automated collection, processing and accessibility of EM data.’, Current opinion in microbiology, 43, pp. 1–8. Available at: 10.1016/j.mib.2017.10.005.

Burt, A. et al. (2021) ‘A flexible framework for multi-particle refinement in cryo-electron tomography.’, PLoS biology, 19(8), p. e3001319. Available at: 10.1371/journal.pbio.3001319.

Carragher, B. et al. (2000) ‘Leginon: an automated system for acquisition of images from vitreous ice specimens.’, Journal of structural biology, 132(1), pp. 33–45. Available at: 10.1006/jsbi.2000.4314.

Cash, J.N. et al. (2020) ‘High-resolution cryo-EM using beam-image shift at 200 keV.’, IUCrJ, 7(Pt 6), pp. 1179–1187. Available at: 10.1107/S2052252520013482.

Cheng, A. et al. (2021) ‘Leginon: New features and applications.’, Protein science : a publication of the Protein Society, 30(1), pp. 136–150. Available at: 10.1002/pro.3967.

Cheng, A. et al. (2023) ‘Fully automated multi-grid cryoEM screening using Smart Leginon.’, IUCrJ, 10(Pt 1), pp. 77–89. Available at: 10.1107/S2052252522010624.

Diebolder, C.A., Dillard, R.S. and Renault, L. (2021) ‘From Tube to Structure: SPA Cryo-EM Workflow Using Apoferritin as an Example.’, *Methods in molecular biology (Clifton*, N.J.), 2305, pp. 229–256. Available at: 10.1007/978-1-0716-1406-8_12.

Dyer, P.M. et al. (2003) ‘Reconstitution of nucleosome core particles from recombinant histones and DNA.’ Methods in Enzymology, 375, pp. 23–24. Available at: 10.1016/S0076-6879(03)75002-2

Efremov, R.G. and Stroobants, A. (2021) ‘Coma-corrected rapid single-particle cryo-EM data collection on the CRYO ARM 300.’, *Acta crystallographica. Section D*, Structural biology, 77(Pt 5), pp. 555–564. Available at: 10.1107/S2059798321002151.

Emsley, P. and Cowtan, K. (2004) ‘*Coot* : model-building tools for molecular graphics’, Acta Crystallographica Section D Biological Crystallography, 60(12), pp. 2126–2132. Available at: 10.1107/S0907444904019158.

Feathers, J.R., et al. (2021) ‘Experimental evaluation of super-resolution imaging and magnification choice in single-particle cryo-EM.’, J Struct Biol X; 5, p. 100047. Available at: doi: 10.1016/j.yjsbx.2021.100047.

Fréchin, L. et al. (2022) ‘High-resolution cryo-EM performance comparison of two latest-generation cryo electron microscopes on the human ribosome.’, Journal of structural biology, 215(1), p. 107905. Available at: 10.1016/j.jsb.2022.107905.

Greenhough, L.A. et al. (2023) ‘Structure and func6on of the RAD51B– RAD51C–RAD51D– XRCC2 tumour suppressor.’, Nature, 619(7970), pp. 650–657. Available at: 10.1038/s41586-023-06179-1.

Grubb, D.T. and Groves, G.W. (1971) ‘Rate of damage of polymer crystals in the electron microscope: dependence on temperature and beam voltage.’, The Philosophical Magazine: A Journal of Theoretical Experimental and Applied Physics, 24(190), 815–828. Available at: 10.1080/14786437108217051.

Hamdi, F., et al. (2020) ‘2.7 Å cryo-EM structure of vitrified M. musculus H-chain apoferritin from a compact 200 keV cryo-microscope.’, PLOS ONE. Edited by C. San Martin, 15(5), p. e0232540. Available at: 10.1371/journal.pone.0232540.

Herzik, M.A., Wu, M. and Lander, G.C. (2017) ‘Achieving better-than-3-Å resolution by single-particle cryo-EM at 200 keV’, Nature Methods, 14(11), pp. 1075–1078. Available at: 10.1038/nmeth.4461.

Keizer, J. et al. (2021) ‘Falcon 4 performance validation by single event analysis’, Microscopy and Microanalysis, 27(S1), pp. 1338–1339. Available at: 10.1017/S1431927621004980.

Kern, D.M. et al. (2021) ‘Cryo-EM structure of SARS-CoV-2 ORF3a in lipid nanodiscs’, Nature Structural & Molecular Biology, 28(7), pp. 573–582. Available at: 10.1038/s41594-021-00619-0.

Kim, L.Y. et al. (2018) ‘Benchmarking cryo-EM Single Particle Analysis Workflow.’, Frontiers in molecular biosciences, 5, p. 50. Available at: 10.3389/fmolb.2018.00050.

Kimanius, D. et al. (2021) ‘New tools for automated cryo-EM single-particle analysis in RELION-4.0’, Biochemical Journal, 478(24), pp. 4169–4185. Available at: 10.1042/BCJ20210708.

Li, Y. et al. (2020) ‘High-Throughput Cryo-EM Enabled by User-Free Preprocessing Routines’, Structure, 28(7), pp. 858–869.e3. Available at: 10.1016/j.str.2020.03.008.

Liebschner, D. et al. (2019) ‘Macromolecular structure determination using X-rays, neutrons and electrons: recent developments in *Phenix*’, Acta Crystallographica Section D Structural Biology, 75(10), pp. 861–877. Available at: 10.1107/S2059798319011471.

Mastronarde, D.N. (2003) ‘SerialEM: A Program for Automated Tilt Series Acquisition on Tecnai Microscopes Using Prediction of Specimen Position’, Microscopy and Microanalysis, 9(S02), pp. 1182–1183. Available at: 10.1017/S1431927603445911.

Mastronarde, D.N. (2005) ‘Automated electron microscope tomography using robust prediction of specimen movements.’, Journal of structural biology, 152(1), pp. 36–51. Available at: 10.1016/j.jsb.2005.07.007.

Merk, A. et al. (2020) ‘1.8 Å resolution structure of β-galactosidase with a 200 kV CRYO ARM electron microscope’, IUCrJ, 7(4), pp. 639–643. Available at: 10.1107/S2052252520006855.

Nakane, T. et al. (2020) ‘Single-particle cryo-EM at atomic resolution.’, Nature, 587(7832), pp. 152–156. Available at: 10.1038/s41586-020-2829-0.

Peck, J.V., Fay, J.F. and Strauss, J.D. (2022) ‘High-speed high-resolution data collection on a 200 keV cryo-TEM.’, IUCrJ, 9(Pt 2), pp. 243–252. Available at: 10.1107/S2052252522000069.

Pettersen, E.F. et al. (2021) ‘UCSF CHIMERAX : Structure visualization for researchers, educators, and developers’, Protein Science, 30(1), pp. 70–82. Available at: 10.1002/pro.3943.

Potter, C.S. et al. (1999) ‘Leginon: a system for fully automated acquisition of 1000 electron micrographs a day.’, Ultramicroscopy, 77(3–4), pp. 153–161. Available at: 10.1016/s0304-3991(99)00043-1.

Punjani, A. et al. (2017) ‘cryoSPARC: algorithms for rapid unsupervised cryo-EM structure determination’, Nature Methods, 14(3), pp. 290–296. Available at: 10.1038/nmeth.4169.

Rawal, Y., et al. (2023) ‘Structural insights into BCDX2 complex function in homologous recombination.’ Nature. 619(7970), pp. 640–649. Available at: doi: 10.1038/s41586-023-06219-w.

Renaud, J.-P. et al. (2018) ‘Cryo-EM in drug discovery: achievements, limitations and prospects.’, Nature reviews. Drug discovery, 17(7), pp. 471–492. Available at: 10.1038/nrd.2018.77.

Robertson, M.J., Meyerowitz, J.G. and Skiniotis, G. (2022) ‘Drug discovery in the era of cryo-electron microscopy.’, Trends in biochemical sciences, 47(2), pp. 124–135. Available at: 10.1016/j.tibs.2021.06.008.

Routine Collection of High-Resolution cryo-EM Datasets Using 200 KV Transmission Electron Microscope. (2022). United States. Available at: 10.3791/63519.

Sader, K. et al. (2020) ‘Industrial cryo-EM facility setup and management’, Acta Crystallographica Section D Structural Biology, 76(4), pp. 313–325. Available at: 10.1107/S2059798320002223.

Stagg, S.M. and Mendez, J.H. (2018) ‘Processing apoferritin with the Appion pipeline.’, Journal of structural biology, 204(1), pp. 85–89. Available at: 10.1016/j.jsb.2018.06.009.

Wieferig, J.-P., Mills, D.J. and Kühlbrandt, W. (2021) ‘Devitrification reduces beam-induced movement in cryo-EM’, IUCrJ, 8(2), pp. 186–194. Available at: 10.1107/S2052252520016243.

Wu, M., Lander, G.C. and Herzik, M.A. (2020) ‘Sub-2 Angstrom resolution structure determination using single-particle cryo-EM at 200 keV’, Journal of Structural Biology: X, 4, p. 100020. Available at: 10.1016/j.yjsbx.2020.100020.

Ye, G., Liu, B. and Li, F. (2022) ‘Cryo-EM structure of a SARS-CoV-2 omicron spike protein ectodomain’, Nature Communications, 13(1), p. 1214. Available at: 10.1038/s41467-022-28882-9.

Yip, K.M. et al. (2020) ‘Atomic-resolution protein structure determination by cryo-EM’, Nature, 587(7832), pp. 157–161. Available at: 10.1038/s41586-020-2833-4.

Zhang, K. et al. (2020) ‘Resolving individual atoms of protein complex by cryo-electron microscopy’, Cell Research, 30(12), pp. 1136–1139. Available at: 10.1038/s41422-020-00432-2.

Zhang, X. et al. (2021) ‘Evolving cryo-EM structural approaches for GPCR drug discovery.’, Structure (London, England: 1993), 29(9), pp. 963–974.e6. Available at: 10.1016/j.str.2021.04.008.

Zhang, Z. et al. (2021) ‘Improving particle quality in cryo-EM analysis using a PEGylation method’, Structure, 29(10), pp. 1192–1199.e4. Available at: 10.1016/j.str.2021.05.004.

